# Trait prediction through computational intelligence and machine learning applied to soybean (*Glycine max*) breeding in shaded environments

**DOI:** 10.1101/2024.01.31.578252

**Authors:** Antônio Carlos da Silva Júnior, Weverton Gomes da Costa, Amanda Gonçalves Guimarães, Waldênia de Melo Moura, Leonardo Lopes Bhering, Cosme Damião Cruz, Renata Oliveira Batista, Jose Barbosa dos Santos, Wellington Ferreira Campos, Anderson Barbosa Evaristo

**Author notes:** **Corresponding author**: Anderson Barbosa Evaristo.

## Abstract

This study aims to identify more relevant predictors traits, considering different prediction approaches in soybean under different shading levels in the field, using methodologies based on artificial intelligence and machine learning. The experiments were carried out under different shading levels in a greenhouse and in the field, using sixteen cultivars. We have evaluated grain yield, which was used as a response trait, and 22 other attributes as explanatory traits. Three levels of shading were used to restrict photosynthetically active radiation (RPAR): 0%, 25%, and 48%. At full sun level (0% RPAR), the traits that presented better predictive performances using a multilayer perceptron were specific leaf area, plant height and number of pods. In the three levels of shading, the plant height trait exhibited the best performance for the radial base function network. Plant height showed the best predictive efficiency for grain yield at 25% and 48% RPAR, for all machine learning methodologies. Computational intelligence and machine learning methodologies have proven to be efficient in predicting soybean grain yield, regardless of shading level.

## 1. Introduction

Soybean [*Glycine max* (L.) Merril] is the fourth most widely grown crop in the world and is one of the prominent species in the global context of agricultural commodities (Ludke et al., 2019; Wiggins et al., 2019; Ravelombola et al., 2020; Costa et al., 2022). Soybean presents great socio-economic importance for Brazil, since it is the basis for various food products and animal feed (Gonçalves et al., 2020). Brazil has consolidated its position as the world’s main soybean producer and exporter. Combined with the USA, both countries account for around 85% of the global soybean trade (USDA, 2020). Brazil’s advanced soybean production system and global relevance are mostly the result of plant breeding.

In this context, there has been increased demand for grain production models that use less input and are more resilient to climate change. Recently, several studies have demonstrated that soybean has been used in alternative cropping systems, such as maize-soybean strip intercropping, agroforestry, tree-based intercropping, and alley cropping systems (Raza et al., 2019; Gagné et al., 2022; Mantino 2019). The obstacle lies in the choice of genetic materials for soybean. The shade provided by a different crop or by the presence of trees induces physiological and/or morphology changes in soybeans (Cheng et al., 2022; Avelino et al., 2023). This influences productivity and quality performance due to a significant reduction of photosynthetically active radiation (Werner et al., 2017; Yang et al., 2014).

One of the major challenges of soybean breeding programs is to predict the importance of traits. Estimating them allows faster progress, selection, and prediction of traits with low heritability and/or measurement difficulties. Although the simultaneous assessment of traits provides a wide range of information, identifying which predictor trait yields the best result is a challenge for the breeder (Parmley et al., 2019). In this context, artificial neural networks (ANNs), decision trees (DT) and their refinements, boosting (BO), random forests (RF), and bagging (BA), present strong potential because they can capture nonlinear relationships between traits (Sousa et al., 2020).

The estimation of the importance of a trait can be performed by artificial neural networks (ANNs) through algorithms such as the one created by Goh (1995), who proposed a modification to Garson’s (1991) algorithm, which consists of partitioning the neural network connection weights to determine the relative importance of each input trait in the network. These methodologies allow for good predictions, determining the importance of a trait to be obtained through measures based, for example, on the Gini and Entropy indexes (Hastie, 2009). These methodologies allow the researcher to quantify the impact of the disruption or the disturbance of the input information to estimate the determination coefficient.

These methodologies have been successfully used in prediction studies. Parmley et al. (2019) evaluated the high-dimensional phenotypic traits in soybean through the machine-learning approach, to predict seed yield for the prescriptive development of cultivars for agricultural practices. Skawsang et al. (2019) applied said methodologies to predict the pest insect population, using the host plant’s climatic and phenological factors. However, there are few reports in the literature related to yield prediction and the importance of traits for grain yield in soybean culture.

Given the above, this work aims to: (1) predict grain yield in soybean under different levels of shading in the field, using methodologies based on artificial intelligence and machine learning; (2) identify more relevant predictors, considering different prediction approaches in soybean.

## 2. Materials and Methods

### 2.1 Experimental data

The experiments were carried out in the Institute of Agricultural Sciences of the Federal University of Vales do Jequitinhonha e Mucuri (UFVJM), Unaí-MG, Brazil (16°26’28.30″ S, 46°54’6.15’’ W, 634 m) between October 2019 and February 2020. Sixteen commercial soybean cultivars (genotypes), 15 of which are of indeterminate growth habit (NS7667 IPRO, NS7780 IPRO, NS7901 RR, RK8115 IPRO, M7110 IPRO, 8579RSF IPRO, RK 6719 IPRO, CD 2728 IPRO, RK 6316 IPRO, CZ37B43 IPRO, 8473RSF RR, 74177RSF IPRO, RK 7518 IPRO, M 6210 IPRO, AS 3680 IPRO) and one of determinate growth habit (NS 8338).

The experiments were conducted at different shading levels in a greenhouse and in the field. In both experiments, three levels of shading were employed: full sun, or 0% – meaning incidence of 100% photosynthetically active radiation (PAR) and restriction of solar radiation on the plants–, 25%, and 48% restriction of PAR (RPAR). All cultivars were grown in the greenhouse and in the field at all PAR restriction levels.

To provide the desired PAR reduction level, a portion of the plants was grown in a greenhouse and in the field under black shade nets that allowed 18% of the incident light to pass through, thus providing 25% PAR reduction. Meanwhile, a different portion of the plants was grown in a greenhouse and in the field under black shade nets that allowed 30% of the light to pass through, thus providing 48% PAR reduction. To determine PAR reduction, the photosynthetic photon flux density (PPFD) was measured throughout the day using a PAR meter (Apogee quantum meters, model MQ-200), during the entire duration of the experiment. Average PAR reduction values were determined and compared with measurements made under full light conditions (i.e., no shade). In the greenhouse, the experiments were conducted between September 2020 and January 2021, and the soybean plants were grown in 10 L plastic pots. The substrate was composed of 50% soil (69.5% clay content) and 50% sand mixed with 36 g of dolomitic limestone, 8 g of simple superphosphate, and 1.60 g of potassium chloride per pot. The experiment was designed in randomized blocks with three replications. The experimental unit consisted of plastic pots, with one plant per pot.

In the field (16°26’2.27’’S, 46°55’28.99’’ W), the experiments were conducted during the rainy season and without supplementary irrigation. Sowing was carried out in October 2020, and harvest took place in March 2021. Each experimental plot consisted of four 6-meter-long rows of plants, with 0.5m of space between the rows, per cultivar. Fertilization and phytosanitary management were carried out according to the recommendations for soybean crops. After 20 days of plant emergence, the shading was installed in the field. The shade nets were installed 1.6m above ground level, so as not to harm soybean development. In the field, the experiment was designed in randomized blocks with three replications.

### 2.2 Evaluated traits

#### Greenhouse

When each genotype reached stage R6 (full seed), the following traits were measured:

i) Leaflet area (LA, cm²), determined with a leaf area meter (LI-COR, model LI-3100), using the central leaflet of the third leaf from the apex;
ii) Chlorophyll a (cl*a*), chlorophyll b (cl*b*) and carotenoids (car), using leaf discs obtained from the central leaflet of the third leaf from the apex, with the aid of a manual hole punch, with 8.6 mm in diameter, with quantified absorbances of 480, 649 and 665 nm measured in a Molecular Devices spectrophotometer and the calculations by Wellburn (1994) expressed as chlorophylls in mg of chlorophyll per g of dry mass of leaf tissue. iii)Chlorophyll fluorescence, determined by maximum photosystem II quantum efficiency (Fv/Fm), maximum fluorescence (MVF) and non-photochemical dissipation (NPQ), using a pulse-modulated fluorometer (Junior-PAM, Germany). The leaves were exposed to a weak pulse of far-red light (1-2 µmol photons m^-2^s^-1^) to determine initial fluorescence (F0). Then, a pulse of saturating light, with an irradiance of 6000 µmol (photons) m^-2^s^-1^ and duration of 1s, was applied to estimate the maximum fluorescence (MVF).
iii) SPAD index, obtained through a portable digital chlorophyll meter (Falker brand, chlorofilog model) by measuring the central leaflet of the third leaf from the apex to the base.
iv) Specific leaf area (SLA), obtained by removing 5 leaf discs from the leaflet center of the third leaf from the apex to the base, which were dried in a forced circulation oven at 70°C for 48 hours, after which the dry mass was determined on an analytical balance.
v) Canopy temperature average values (CTA), obtained through images of the plants captured with an infrared thermal camera (model E5, FLIR), using 10 points to determine the average, where each pixel generates a temperature value.
vi) Light extinction coefficient (LEC), obtained by the quotient between the amount of PAR that reaches the apex of the plant and the amount of PAR near the ground in. A PAR meter (Apogee quantum meters, model MQ-200) was used to quantify the PAR present in each position of the plant.

When the plants reached full flowering, stage R2 (full flowering, with an open flower at one of the two uppermost nodes), the days for flowering (DF) were determined. At stage R8 (full maturity, when 95% of the pods have reached their mature color) the following were evaluated:

i) Plant height (PH, cm), measured from the ground to the plant apex.
ii) Hypocotyl diameter (D, mm), determined using a digital caliper, close to ground level.
iii) Number of branches (NB) per plant.
iv) Node number (NN) on the main stem.
v) Inter-node length (IL, cm), obtained by dividing the length of the stem by the number of the node in the main stem.
vi) Yield components: the number of pods per plant (NPP), the number of seeds per pod (NSP) from a sample of 30 pods per plant, and the hundred seed weight (HSW, g), determined by counting three subsamples of grain.

The grain moisture content was determined using a digital moisture meter and the grain mass was quantified in an analytical balance. HSW data were adjusted to 13% grain-moisture content.

#### Field

In the field experiment, approximately after eight days from the beginning of stage R8, the two 5-meter-long central rows were manually harvested, discarding 0.5m at the beginning and end of the plot. These plants were threshed using a stationary threshing machine. The grains were weighed and the moisture was determined using a portable grain moisture meter (model G650i, Gehaka brand). Grain moisture was adjusted to 13% moisture and yield was determined in kg ha^-1^. The trait grain yield (GY, Kg ha^-1^).

### 2.3 Training and Validation Sets

For better network efficiency, before training and validation, the data were normalized in the interval between −1 and 1 for all methodologies. The training data set, in each location, was established by 2/3 of the phenotypic information, using the strategy of aggregating information from two of the three repetitions for training, and the information from the other repetition was used as a validation set. In this cross-validation strategy, individuals from each repetition participated at mean once in the validation data set in cross-validation (k-fold) k = 3 partitions. Other studies also utilized this strategy (González-Camacho et al., 2012).

### 2.4 Multilayer Perceptron - MLP

The maximum number of training seasons was set at 5,000; the mean square error (MSE), as a criterion to stop processing the network, was defined as 1.0 × 10^−3^. All trained networks had a neuron in the output layer and a single hidden layer, with 10 neurons. The sigmoid tangent activation function was used in the hidden layer. The training algorithm was employed was *Bayesian regulation backpropagation*, to quantify the efficiency of the prediction (*R*^2^).

The importance of predictors through the MLP network was quantified using two techniques. The first one, based on Garson’s (1991) algorithm modified by Goh (1995), consists of partitioning the neural network connection weights to determine the relative importance of each input trait within the network (Silva Junior et al. 2021).

The equation of the relative importance of the trait is equal to:

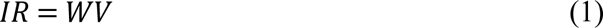

The matrixial model is shown below:

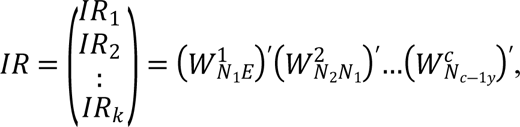

where, 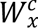 represents the matrix of weights of the layer c neuron, considering *N_j_* neurons and *N_j-1_* inputs; *E* is the first neuron that starts from inputs; *y* refers to the desired output layer, and *IR* is the relative importance of the trait.

After network stabilization, the importance of traits (inputs) was obtained, considering the impact of de-structuring or disturbing the information of a given input to estimate the coefficient of determination (Silva Junior et al., 2021).

The second technique was based on the relative importance of the trait by the permutation of *R*^2^, which is described in the following equation:

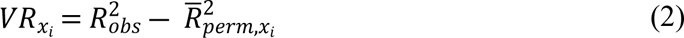

where, 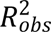 is the *R*^2^ of the RNA model adjusted to the observed predictor and response traint; 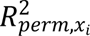, is the *R*^2^ of the ANN model fitted to the modified dataset where *x*_*i*_ is permuted; 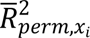: is the average value of 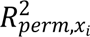 after m^th^ permutation of the datasets.

### 2.5 Radial Base Function Network – RBF

The radial base function network is characterized by having only one hidden layer and making use of the Gaussian activation function (Cruz & Nascimento 2018). To better predict grain yield, the structure of the RBF was established with 10 to 30 neurons (increased by 2, with each processing), and the radius was established between 5 and 15 (increased by 0.5). The efficiency of the prediction was measured by the R^2^, and the relative importance of each entry was measured by the technique of destroying the information of each explanatory trait, as already described for MLP.

### 2.6 Machine learning for the soybean yield prediction

To establish soybean yield prediction through a machine learning approach, the decision tree and its refinements, random forest, bagging, and boosting were employed. The soybean yield prediction (PRY) is described in the following Equation:

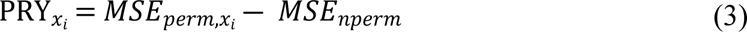

where, *MSE_perm,x_i__* is the permutation of the values of each trait in the dataset where *x*_*i*_ is swapped, and *MSE_nperm_* represents the values of the estimate of the original non-permuted variable data.

All the procedures of data analyses that were used to obtain statistics related to the parameters of interest were processed in the R platform (R Core Team et al., 2022), software Genes (Cruz, 2016), with the use of an interface with MATLAB software (Matlab, 2011).

## 3. Results

### Prediction by different approaches

The R² of the estimates for all methodologies, using explanatory traits for soybean grain yield in the field and the auxiliaries evaluated in a greenhouse, are shown in Figure 1. Among the computational intelligence and machine learning methodologies, RF presented the highest estimate of the maximum coefficient of determination (R²) for the environment without shade (0% RPAR). The ANNs methodologies obtained better results in shaded environments (Figure 1). RBF technique obtained better R² for the 25% RPAR environment, while MLP presented better results in the 48% RPAR environment.

**Figure 1.**
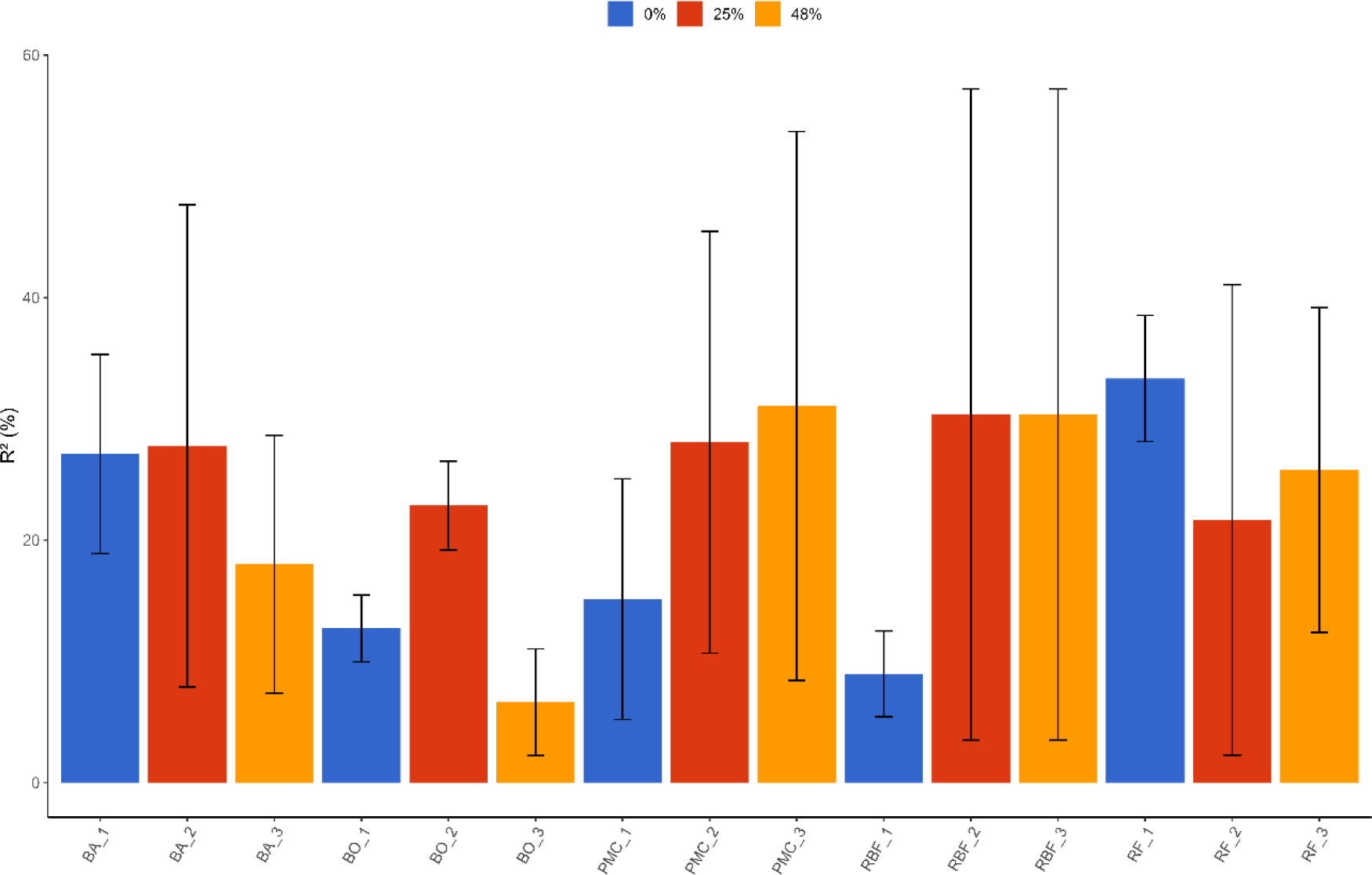
Maximum estimate of the coefficient of determination in three shading levels to predict grain yield (GY) in soybean; MLP: multilayer perceptron; RBF: radial base network; RF: *random forest*; BA: *bagging*; BO: *boosting*. The numbers in the methodologies represent the levels of shading. 1: SL 0% (100% PAR); 2: SL 25% RPAR, and 3: SL: 48% RPAR. Vertical bars represent standard deviation.

The relative contributions estimated by the weight of connections using the RFB and MLP are shown in Figure 2. At 0% RPAR level, which corresponds to the full-sun environment, the traits that presented the best predictive performances were SLA, D, and MVF, respectively, using the MLP network. Regarding the 25% RPAR and 48% RPAR shading levels, CTA and NB obtained the highest relative contributions by the weight of connections through the MLP.

**Figure 2.**
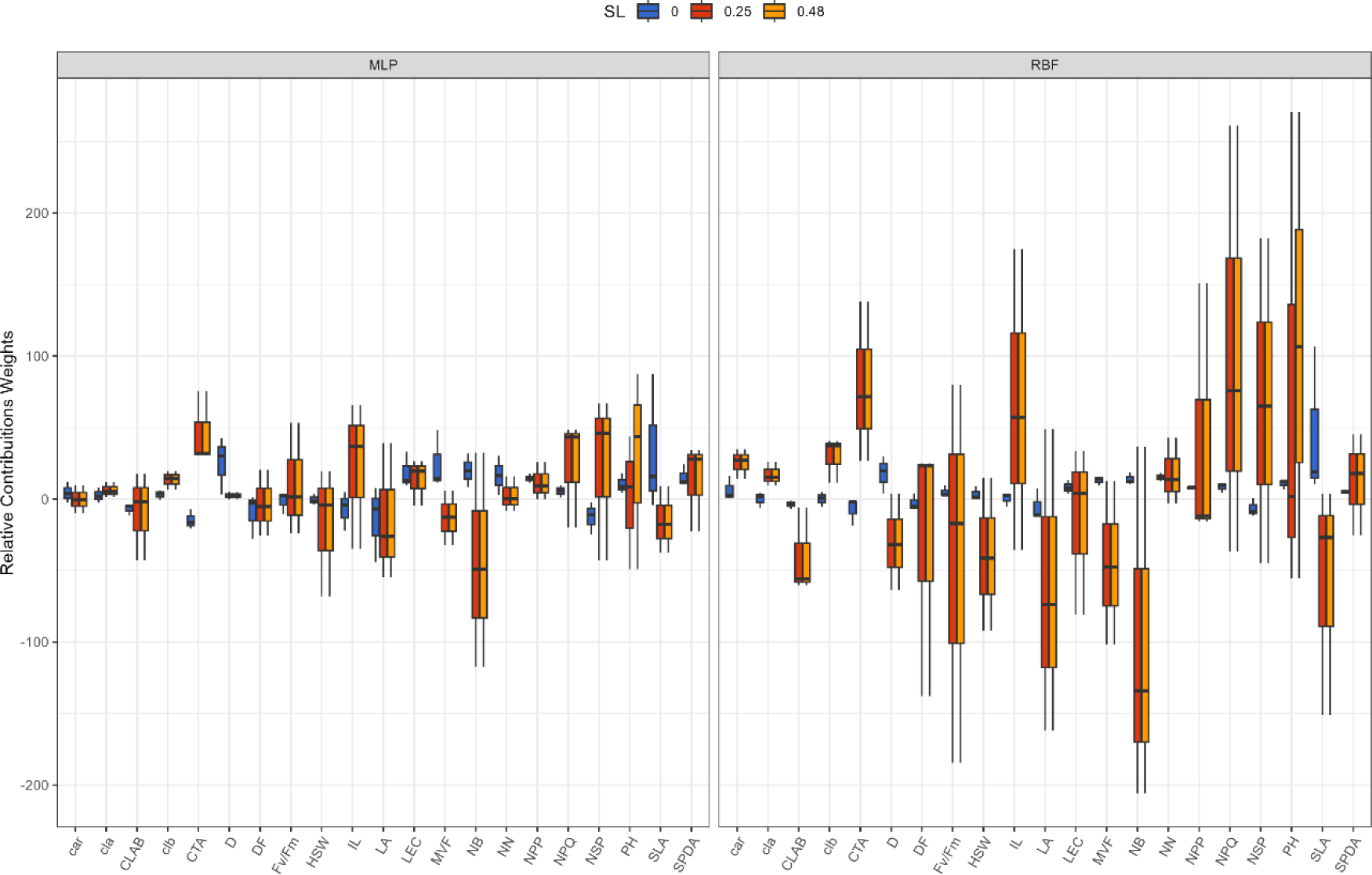
Relative connection weight estimated by Garson’s method (1991), modified by Goh (1995), of 21 traits to predict soybean grain yield in the evaluated field in pots under three shading levels. SL 0%RPAR (100% PAR); SL 25%RPAR and SL: 48%RPAR; LA= area leaf; CTA: canopy temperature average values; SPDA: index of chlorophyll meter; SLA: Specific leaf area; PH: plant height; D: diameter; cl*a*: Chlorophyll A; cl*b*: Chlorophyll B; car: carotenoids; CLAB: Chlorophyll A/B ratio; Fv/Fm: maximum photosystem II quantum efficiency; FVM: maximum fluorescence; NPQ: non-photochemical dissipation; IL – inter-node length; LEC: light extinction coefficient; DF: days for flowering; NN: Number of nodes; NPP: Number of pods per plant; NB: number of branches; NPM: number of pods/m²; NSP: number of seeds per pod; Hundred seed weight (HSW).

For the RBF network, the 0% RPAR SLA presented the best performance, followed by D, NN, NB, MVF, and PH, respectively. In the two shading levels (25% and 48% RPAR), the traits that showed the best performance were NPQ, CTA, and PH. In the module, NB also showed a high relative contribution.

Figure 3 describes the quantification of the importance of soybean traits by randomizing the information of an input trait after establishing the MLP. In this Figure, the values are used after causing disturbances in the input traits with the action of assigning a zero value to the trait in each explanatory trait. For this method, the traits that obtained the lowest estimate of *R*^2^ were D, LEC, MVF, and NN at 0% RPAR level. All traits showed similar behavior at 25% and 48% RPAR levels. The traits that showed the best performance to predict productivity in soybean were DF, D, NN, SLA, CLAB, HSW, CAR, CLA, LEC, and NSP.

**Figure 3.**
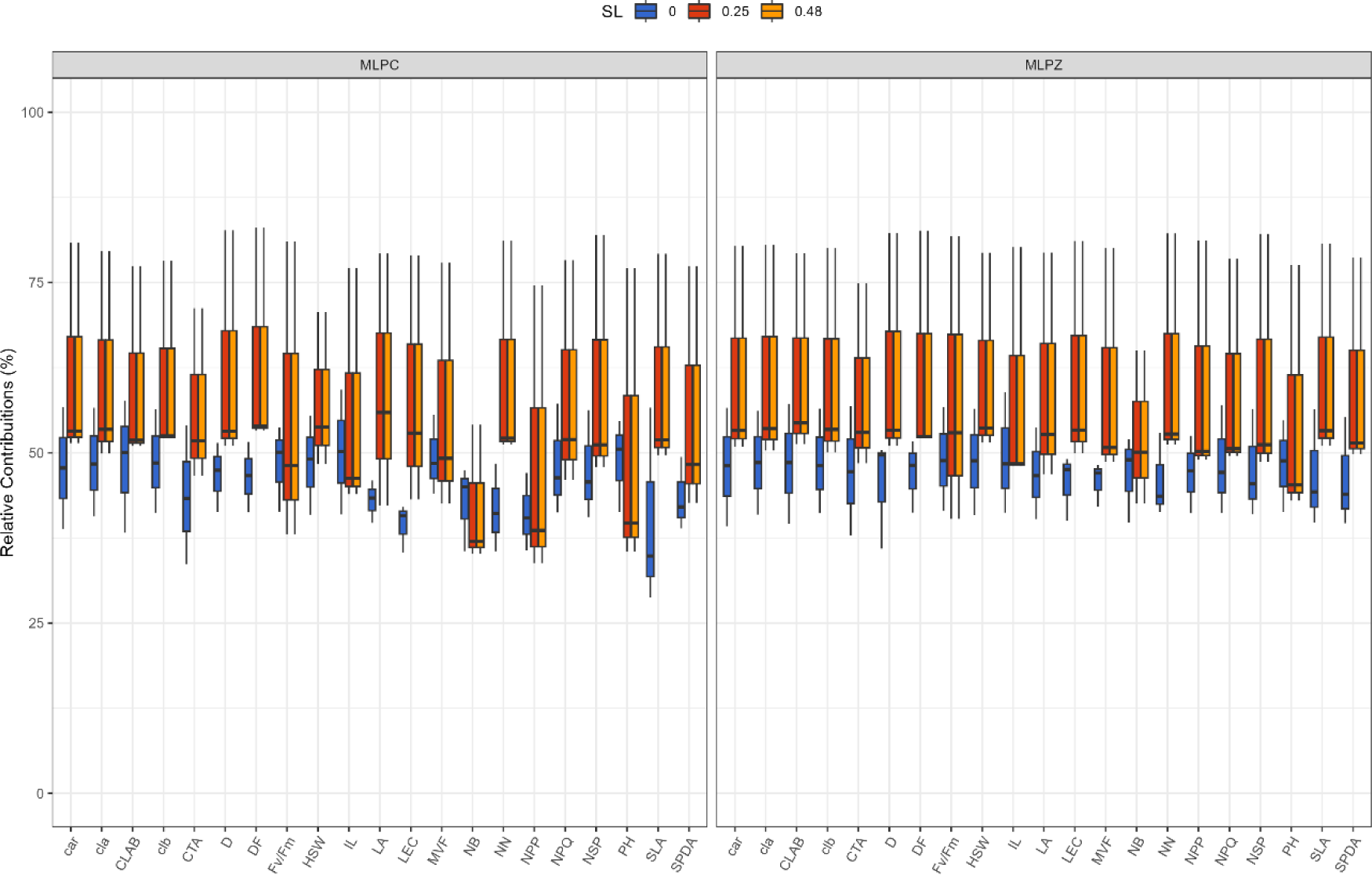
Estimates of the coefficient of determination for grain yield prediction in soybean using MLP assigning zero value and randomizing genotype information. MLPZ: assigning value zero; MLPC: randomizing genotype information. Three shading levels: SL 0%RPAR (100% PAR); SL 25%RPAR and SL: 48%RPAR; LA: area leaf; CTA: canopy temperature average values; SPDA: index of chlorophyll meter; SLA: Specific leaf area; PH: plant height; D: diameter; cl*a*: Chlorophyll A; cl*b*: Chlorophyll B; car: carotenoids; CLAB: Chlorophyll A/B ratio; Fv/Fm: maximum photosystem II quantum efficiency; FVM: maximum fluorescence; NPQ: non-photochemical dissipation; IL: inter-node length; LEC: light extinction coefficient; DF: days for flowering; NN: Number of nodes; NPP: Number of pods per plant; NB: number of branches; NPM: number of pods/m²; NSP: number of seeds per pod; Hundred seed weight (HSW).

Figure 4 shows the results of quantifying the importance of phenotypic traits using RBF after trait permutation and assigning a zero value to input traits. Using this methodology, it was observed that car, HSW, clb, cla, and IL were the traits that were most important in predicting soybean yield at 0% RPAR. At shading levels 25% RPAR and 48% RPAR, in addition to the parameters mentioned above, D, DF, NPP, SLA, and NN also presented high values for relative contributions.

**Figure 4.**
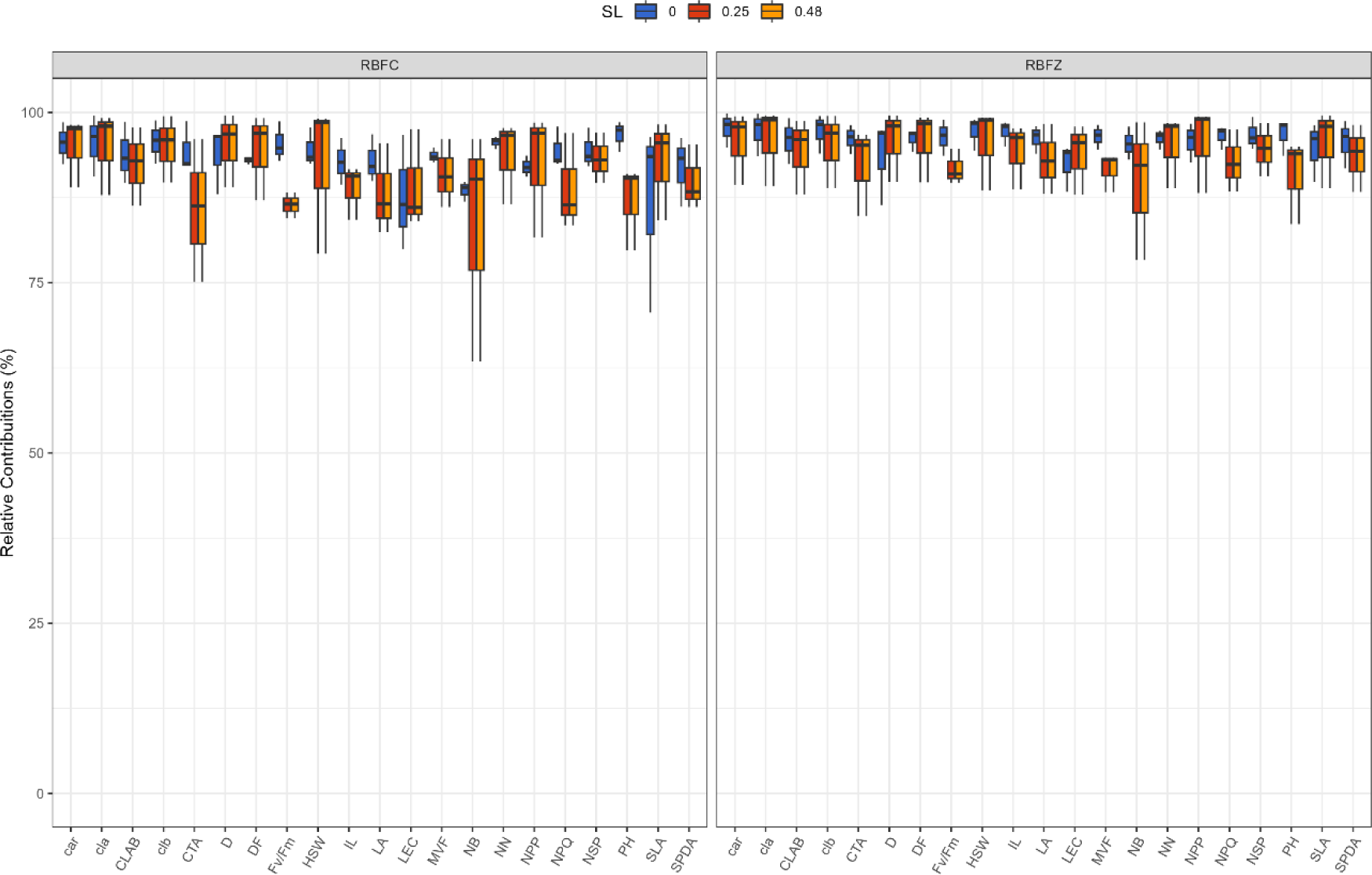
Estimates of the coefficient of determination for grain yield prediction in soybean using the RBF assigning zero value. RBFZ: assigning zero value; RBFC: randomizing genotype information. Three shading levels; SL 0%RPAR (100% PAR); SL 25%RPAR and SL: 48%RPAR; LA: area leaf; CTA: canopy temperature average values; SPDA: index of chlorophyll meter; SLA: Specific leaf area; PH: plant height; D: diameter; cl*a*: Chlorophyll A; cl*b*: Chlorophyll B; car: carotenoids; CLAB: Chlorophyll A/B ratio; Fv/Fm: maximum photosystem II quantum efficiency; FVM: maximum fluorescence; NPQ: non-photochemical dissipation; IL: inter-node length; LEC: light extinction coefficient; DF: days for flowering; NN: Number of nodes; NPP: Number of pods per plant; NB: number of branches; NPM: number of pods/m²; NSP: number of seeds per pod; Hundred seed weight (HSW).

Contrarily, after the randomization of their values, the traits that showed the best performance to predict productivity in soybean were PH, D, DF, clb, NN, cla, and car at all RPAR levels, with the exception of D and DF at 0% RPAR, as well as PH at 25% RPAR and 48% RPAR. The traits that showed the lowest mean estimate of the coefficient of determination were SLA and NB.

The averages of the relative contributions of the explanatory traits to the prediction of soybean grain yield are shown in Table 1. This prediction is obtained by estimating the mean squared error increment percentage (MSEI), which is constructed by changing the values of each trait in the data set and comparing them with the set prediction of the trait’s original non-permuted data. Similar to the strategy used for the computational intelligence methodologies of the MLP and RBF networks (for which greater values indicated greater importance of this trait’s *R*^2^ in the model), in the machine learning approach, the importance of the explanatory trait is related to the estimate of the average decrease in model accuracy through the MSEI. Therefore, a greater estimate means greater importance of the trait.

**Table 1.** Means of the relative contributions of the explanatory variables for the prediction of soybean grain yield by estimating the minimum squared error increment percentage (MSEI)

| Trait | 0% |  |  | 25% |  |  | 48% |  |  |
| --- | --- | --- | --- | --- | --- | --- | --- | --- | --- |
|  | BO | BA | RF | BO | BA | RF | BO | BA | RF |
| IL | **6.03** | -0.13 | 1.66 | 5.67 | **3.10** | **5.34** | 4.67 | **4.79** | **3.84** |
| LA | 4.71 | 0.26 | -0.58 | **6.08** | -0.86 | 2.44 | **5.08** | -1.08 | 0.81 |
| SLA | **7.47** | -2.59 | -0.76 | **6.72** | -1.65 | -0.22 | **6.90** | -1.58 | -2.17 |
| PH | 4.02 | -0.06 | **2.19** | **6.35** | **4.59** | **3.91** | **6.95** | **5.08** | **4.87** |
| car | 5.74 | -0.35 | 0.64 | 3.77 | 0.12 | -0.78 | 3.79 | -0.27 | -2.16 |
| cla | 3.02 | -0.44 | -2.34 | 4.94 | 0.40 | 3.10 | 4.50 | 0.30 | 1.00 |
| cla/b | 3.29 | 0.76 | -0.12 | 5.47 | -0.61 | 0.52 | 5.90 | 0.22 | 0.16 |
| clb | 2.02 | -2.14 | 0.06 | 4.40 | -1.72 | 1.51 | 4.55 | 0.86 | **2.27** |
| D | 5.43 | **7.11** | **4.18** | 4.64 | -0.79 | -0.42 | 4.88 | 0.13 | 0.62 |
| DF | 0.51 | 0.62 | 0.09 | 0.71 | -0.82 | -0.32 | 0.96 | 0.88 | 1.70 |
| LEC | 3.05 | 0.07 | **2.62** | 1.96 | -1.13 | -0.22 | 1.99 | -0.84 | -0.54 |
| Fv/Fm | 4.53 | 2.86 | -2.17 | **7.40** | 2.08 | 1.58 | **7.25** | 1.57 | 2.05 |
| MVF | **6.94** | **9.66** | **4.77** | 3.04 | 0.28 | -0.69 | 3.67 | 3.12 | 0.75 |
| HSW | 4.44 | -1.64 | -3.44 | 4.61 | 1.09 | 2.81 | 4.68 | **6.12** | **2.42** |
| NSP | 3.22 | 1.33 | 1.74 | **6.32** | **3.08** | **3.26** | **7.32** | **4.57** | **5.12** |
| NB | 3.23 | 0.36 | 1.21 | **6.04** | 1.52 | 2.68 | **6.04** | 0.57 | 2.66 |
| NN | 4.43 | 1.47 | 1.11 | 1.55 | -0.34 | -1.73 | 0.55 | -0.20 | -1.99 |
| NPQ | 3.88 | -1.43 | 0.40 | 2.77 | **8.82** | **4.93** | 3.71 | **9.77** | **4.95** |
| NPP | **7.77** | 3.36 | 0.99 | **7.39** | -2.43 | 0.95 | **6.39** | 1.11 | -0.65 |
| SPAD | **8.28** | 1.88 | 0.23 | 3.66 | 1.04 | -1.85 | 2.61 | 0.11 | -0.61 |
| CTA | **7.89** | 1.25 | 0.68 | **6.38** | 0.89 | 1.78 | **5.70** | 3.01 | 0.96 |

RF: random forest; BA: bagging; BO: boosting. The numbers in the methods represent the levels of shading SL 0%RPAR (100% PAR); SL 25%RPAR and SL: 48%RPAR; IL: inter-node length; LA: area leaf; CTA: canopy temperature average values; SPDA: index of chlorophyll meter; SLA: Specific leaf area; PH: plant height; D: diameter; cla: Chlorophyll a; clb: Chlorophyll b; car: carotenoids; cla/b: Chlorophyll A/B ratio; Fv/Fm: maximum photosystem II quantum efficiency; MVF: maximum fluorescence; NPQ: non-photochemical dissipation; LEC: light extinction coefficient; DF: days for flowering; NN: Number of nodes; NPP: Number of pods per plant; NB: number of branches; NPM: number of pods/m<sup>2</sup>; NSP: number of seeds per pod; HSW: Hundred seed weight.

The trait that showed the highest MSEI estimate in all machine learning methodologies was the number of seeds per pod (NSP) and plant height (PH) at shading levels 25% and 48% RPAR. When all the evaluated attributes are considered, this trait has shown relevance in the prediction of soybean grain yield using machine learning approaches. However, NSP did not show the importance of shading level 0% RPAR (100% PAR). At this level, the machine learning methodologies coincided with the quantification of MVF as an important trait for grain yield prediction.

## 4. Discussion

In our study, we compared various approaches to trait selection to identify the most relevant predictive traits for soybean grain yield. This is particularly important because the traits used in our research can be challenging to obtain and to evaluate, especially when dealing with a large number of genotypes, which can be costly and labor-intensive (Ferreira et al., 2015). Machine learning techniques have consistently been recognized as efficient tools to quantify the relative importance of traits due to their simplicity, their ability to work without making assumptions about trait distributions, and their robustness in handling various data complexities (Tan et al., 2014; Beucher et al., 2019; Costa et al., 2020; Silva Junior et al., 2021).

The use of computational intelligence and machine learning techniques to predict soybean grain yield is in line with previous studies that have emphasized the effectiveness of these approaches (Tan et al., 2014; Beucher et al., 2019; Silva Junior et al., 2021). The results indicate that different methodologies are more suitable for different environmental conditions, such as full sun or shading, in agreement with the observation that selecting relevant features varies depending on the growth environment (Zhang et al., 2019; Liu et al., 2018). This research reinforces the idea that machine learning and computational intelligence allow for more precise adaptation to specific growth conditions, since these methodologies also offer the advantage of capturing complex factors in predictive models and are capable of modeling data in a non-linear and non-parametric manner (Osco et al., 2020).

The results obtained by the different methodologies were consistent in pointing out that the characteristics that presented greater relative importance varied according to the environment. Plant diameter was important in all the environments evaluated, while the importance of photosynthetic characteristics was demonstrated in environments under full sun or little shade. In full-sun environments (0% RPAR), traits such as MVF, IL, D, clb, and cla emerged as the strongest predictors. This aligns with the well-documented soybean sensitivity to shading due to their C3 photosynthetic pathway, which requires ample light for efficient photoassimilation (Casaroli et al., 2007). This sensitivity is reflected in the decreased productivity observed under shading conditions (Yao et al., 2017; Balbinot-Junior et al., 2018a, 2018b). Other studies have emphasized the importance of different features, such as diameter, pods per plant, and plant height in shaded environments (Kantolic et al., 2013; Balbinot-Junior et al., 2016).

In shaded environments, in addition to plant diameter, other characteristics stood out as being important in predicting grain yield, such as flowering duration (DF), number of pods per plant (NPP), number of seeds per pod (NSP), plant height (PH), weight of 100 seeds (HSW), carotenoids (car), chlorophyll a (cl*a*), chlorophyll b (cl*b*), luminosity index (IL) and canopy temperature average values (CTA). These characteristics relate to the efficiency in the use of radiation, the plant architecture and the number and weight of the seeds. The observation that the number of seeds per pod (NSP) and plant height (PH) are consistently relevant in shading environments is in line with previous research that has underscored these features as crucial determinants for soybean productivity (Kantolic et al., 2013; Balbinot-Junior et al., 2016).

The method of quantifying feature importance through randomization provides additional insights into the robustness of the identified features. This approach complements previous studies that have emphasized the importance of features such as specific leaf area and flowering duration in different environmental conditions (Zhang et al., 2019; Balbinot-Junior et al., 2016). Furthermore, the variation in feature importance with randomization reflects the adaptability of soybeans to different conditions.

The selection of soybean cultivars with a high number of seeds in shaded environments is essential because, in said environments, low incident radiation can reduce the number of pods produced on most branches of the plant (Kantolic et al., 2013). The increase in plant height in shaded environments is induced by the quality of the light (Schmitt, 1997). The characteristic number of stems favors the development of reproductive structures from which the pods develop and, consequently, it influences production (Ferrari et al., 2018). Teodoro et al. (2015) observed an indirect effect of the number of stems on grain yield due to its association with the number of seeds per plant.

In shaded environments, the importance of traits such as DF, D, NPQ, NSP, IL, car, and cl*a* became evident, depending on the shading level (25% and 48% RPAR). The number of seeds per pod (NSP) and plant height (PH) consistently stood out across all methodologies, underscoring their crucial roles. Low incident radiation in shaded environments can diminish pod production in most branches of soybean plants, making the selection of cultivars with a high seed count essential (Kantolic et al., 2013). The increase in plant height under shading conditions is influenced by light quality (Schmitt, 1997). Furthermore, the number of branches influences the development of reproductive structures, ultimately impacting production (Ferrari et al., 2018; Teodoro et al., 2015).

The implications of this study for soybean selection and breeding in shaded environments are in line with the need to adapt selection strategies based on environmental conditions, as observed in previous studies (Zhang et al., 2019; Balbinot-Junior et al., 2016). The ability to identify key features and their variable importance allows breeders to make informed decisions, contributing to the sustainable improvement of soybean cultivation in diverse environments.

Our findings echo previous research suggesting that soybeans can adapt to low-light environments by optimizing assimilate consumption and light utilization (Hussain et al., 2020). Additionally, selecting soybean varieties with a gradual transition from vegetative to reproductive growth phases can enhance yield, a strategy supported by Fan et al. (2018).

The results of this work advance knowledge on the adaptation of soybeans to shaded environments, which are increasingly common due to integrated planting systems. The results also contribute to the genetic improvement of soybeans, by indicating which characteristics should be prioritized in the selection of genotypes adapted to these environments. The characteristics identified as most important can be used as indirect selection criteria for grain yield, reducing physical effort, cost, use of labor and time spent on trials.

## 5. Conclusion

Computational intelligence and machine learning methodologies were able to quantify the importance of explanatory traits in predicting soybean grain yield.

Differences were found between the variables that are most important for grain yield analysis according to the environment. The diameter of the plants was important in all environments evaluated, while the importance of photosynthetic characteristics was demonstrated in environments under full sun or with little shading.

## Acknowledgments

Thanks to Conselho Nacional de Desenvolvimento Científico e Tecnológico (CNPq) for the financial support, Fundação de Amparo à Pesquisa do Estado de Minas Gerais (FAPEMIG) and the Coordenação de Aperfeiçoamento de Pessoal de Nível Superior – Brasil (CAPES) - Finance Code 001”.

